# Impact of DNA polymerase choice on assessment of bacterial communities by a *Legionella* genus-specific next-generation sequencing approach

**DOI:** 10.1101/247445

**Authors:** Rui P. A. Pereira, Jörg Peplies, Ingrid Brettar, Manfred G. Höfle

## Abstract

The library preparation step is a major source of bias in NGS-based studies. Several PCR-related factors might negatively influence the application of NGS tools in environmental studies and diagnostics. Among the most understudied factors are DNA polymerases. In our study, we evaluated the effect of DNA polymerase type on the characterisation of bacterial communities, more precisely *Legionella*, using a genus-specific NGS approach. The assay with proof-reading high fidelity KAPA HiFi showed better amplification yield than the one with widely used non-proofreading HotStarTaq. *Legionella* community richness metrics were significantly overestimated with HotStarTaq. However, the choice of DNA polymerase did not significantly change the community profiling and composition. These results substantiate the use of proof-reading high fidelity DNA polymerases in NGS assays and highlight the need of considering the impact of different DNA polymerases in comparative studies and future guidelines for NGS-based diagnostic tools.

## INTRODUCTION

Since its arrival, next-generation sequencing (NGS) has undergone fast and continuous progress in terms of technology, allowing enhanced performance, throughput and accuracy in a shorter time [1]. NGS methodologies have been widely applied in environmental microbiology to provide a broader characterisation of bacterial community structure, a better comprehension of the influence of environmental and anthropogenic drivers in both the emergence and persistence of bacterial pathogens; and for the detection and characterisation of clinically and environmentally relevant organisms [2–4]. One of the most environmentally and clinically important water-based bacterial taxa worth studying is *Legionell*a. *Legionella* species are persistently and abundantly found in a highly diverse set of aquatic environments and more than 25 species have been associated with disease [5]. Legionellosis outbreaks are frequently reported and have been mostly linked to hot water settings and foremost cooling towers, suggesting the need of a continuous environmental surveillance. NGS assays have already been demonstrated to have the capacity to provide a detailed understanding of the spatio-temporal dynamics and diversity of *Legionella* species and be used as reliable and precise monitoring tools of pathogenic species, such as *L. pneumophila* [6].

NGS has the potential to be an upgrade to the current accepted methodologies, and consequently be translated into active frameworks for routine surveillance and investigation of man-made freshwater systems. However, thorough *in vitro* and *in situ* validation studies are needed to evaluate if NGS-based methodologies are precise and reliable methodologies that are worth to be widely implemented and applied. Guidelines have already been presented and implemented for the use of NGS in clinical diagnostics and genetics screening [7–9]. Yet, no recommendations, guidelines or regulatory approaches currently exist for validation and establishment of the assays and interpretation of the results in environmental settings. Among the factors to consider in the validation of a NGS assay are the potential bias introduced by library preparation and sequencing steps, which can affect not only sensitivity and specificity of the molecular assay developed but also inter-laboratory comparison studies. The potential of NGS as an accurate and sensitive tool for environmental research and diagnostics can be affected by the choice of the primers, targeted taxonomic group, PCR thermo-cycling conditions, template concentration and DNA polymerase [10–13].

In this study, our aim was to evaluate the impact of the choice of the DNA polymerase have on assessment of microbial diversity, more specifically on the alpha- and beta-diversity of *Legionella* community in freshwater systems, with a genus-specific 16S rDNA-based NGS approach.

## MATERIALS AND METHODS

To study the effect of the DNA polymerase on *Legionella* libraries community diversity metrics and structure, 8 freshwater samples, i.e. 2 cold drinking water (A and B), 2 hot drinking water (C and D) and 4 cooling tower water (E to H) samples, were collected at the campus of the Helmholtz Centre for Infection Research (HZI), in Braunschweig, Germany, and processed as previously described [6,14,15]. These freshwater samples were then analysed by a genus-specific NGS approach using two optimised and independent assays with two hot-start DNA polymerases, HotStarTaq (Qiagen, Hilden, Germany) and proofreading KAPA HiFi (KAPA Biosystems, Wilmington, MA, USA). A 16S rRNA gene fragment, with a length of 421 bp, comprising the V3-V4 hypervariable regions was amplified with *Legionella* genus-specific primer pair Lgsp17F 5´-GGCCTACCAAGGCGACGATCG-3’/ Lgsp28R 5´-CACCGGAAATTCCACTACCCTCTC-3’ [16]. Detailed descriptions of the used Illumina MiSeq-based approach with both DNA polymerases as well as of the 16S rDNA data processing and taxonomic classification by the bioinformatics pipeline of SILVA project are given elsewhere [17,18].

## RESULTS

Before evaluating the effect of the DNA polymerase on community diversity metrics, we compared the yield of the two optimised assays by equivolume pooling of all sample libraries and sequencing in a single Illumina MiSeq 250 bp paired-end run. The data revealed a statistically significant difference between the sample sets concerning the number of retrieved 16S rRNA *Legionella* gene sequences per sample, after quality filtering and taxonomic assignment (Welch’s t-test, P<0.05). The mean number of *Legionella* 16S rRNA gene sequences per sample in the sample set amplified by KAPA HiFi was significantly higher (52,385 ± 12,907) than in the one amplified by HotStarTaq (37,343 ± 8,995). Of the 8 samples, 7 had a higher number of sequences when amplified with KAPA HiFi, representing, on average, an increase of 45% when compared with HotStarTaq. This discrepancy does not seem to be an outcome of poor sequence quality as the sample sets presented a similar percentage of rejected sequences during quality filtering (1%). These results suggest that the optimised assay with KAPA HiFi polymerase provided a higher *Legionella* amplification yield than the one with HotStarTaq polymerase, regardless of the observed inferior specificity to *Legionella* genus (86% ± 11% against 97% ± 2%). Nonetheless, rarefaction analyses revealed Operational Taxonomic Units (OTU) values levelling off markedly for all samples (data not shown).

Alpha-diversity metrics, i.e. observed OTU richness (OTUobs) and Shannon’s diversity index (*H’*), were calculated, using the software package Explicet [19], for the *Legionella* community of the 8 water samples amplified with both KAPA HiFi and HotStarTaq DNA polymerases, after clustering of NGS reads into OTUs using an identity criterion of 98%. (**Fig. 1**). The use of HotStarTaq, comparatively to KAPA HiFi, showed a steeper slope and exhibited a higher *Legionella* richness with a statistically significant increase of the number of OTUs observed (Welch’s t-test, p<0.05) regardless of the sample (Fig. 1(a)). The sample set when amplified by HotStarTaq had an averaged observed richness of 2389 ± 851 OTUs. However, when amplification performed with KAPA HiFi, the sample set showed an averaged observed richness of 1846 ± 469 OTUs, representing a decrease of 23% to the values of *Legionella* observed richness shown with HotStarTaq. Also, the number of singletons and doubletons were reduced by an average of 20% and 22%, respectively, with KAPA HiFi. Similar findings were observed when separately amplifying technical replicates (n=3) of a single strain (*L. pneumophila* ATCC 33152^T^), with a total reduction of 38% in singletons and doubletons detected when using KAPA HiFi (data not shown).

**Fig. 1.**
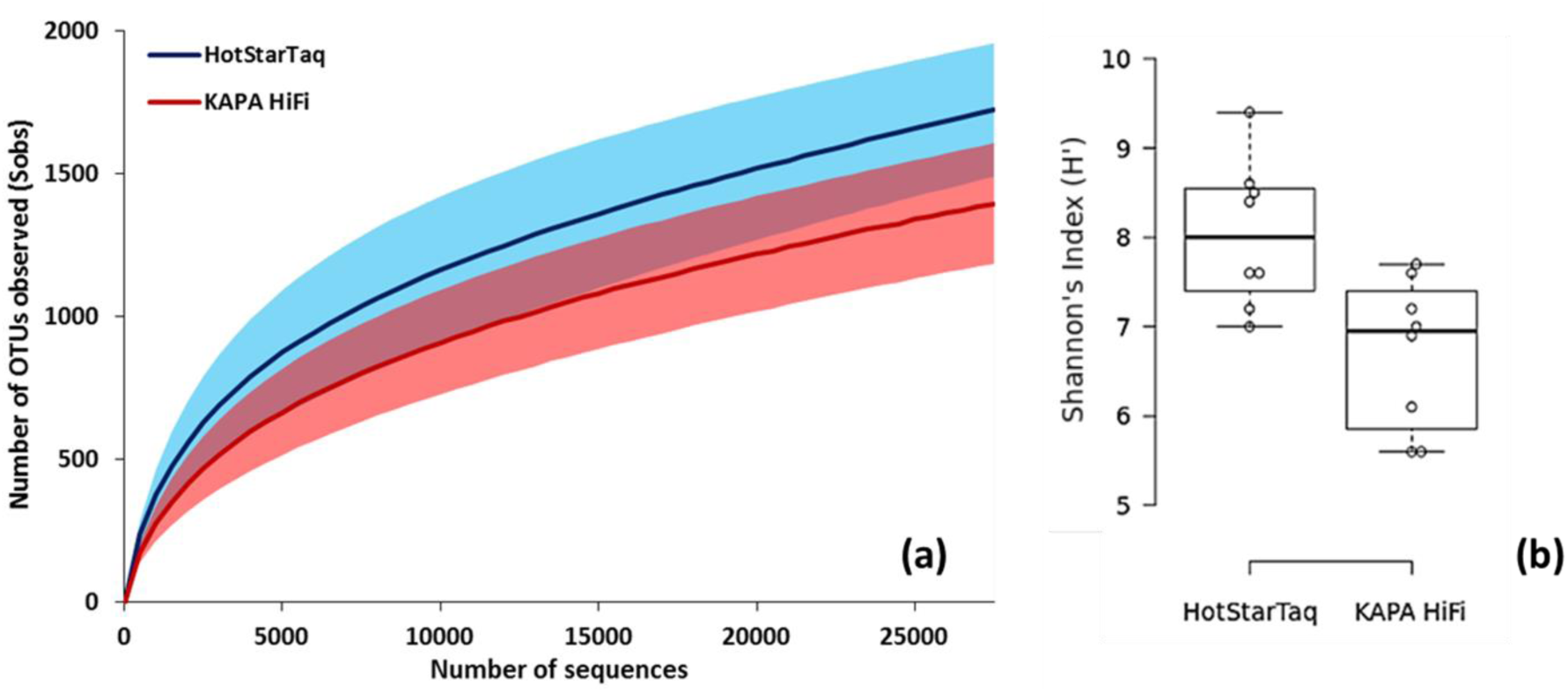
Alpha-diversity assessment of *Legionella* communities in 8 water samples with HotStarTaq and KAPA HiFi DNA polymerases. **(a)** Rarefaction curves showing progress of number of OTUs observed with increasing number of sequences analysed. Number of OTUs observed were calculated after normalisation to the sample with the lowest number of sequences. Lines indicate mean values with corresponding coloured shaded standard deviation for HotStarTaq DNA polymerase (blue) and KAPA HiFi DNA polymerase (red). (**b**) Boxplots of Shannon’s diversity index (*H’*) of 8 water samples amplified with both DNA polymerases. Bars within boxes indicate the median value of diversity. Whiskers extend to data points that are less than 1.5 × the interquartile range away from 1^st^/3^rd^ quartile.

Moreover, the comparison of the two sample sets showed a statistically significant difference in the Shannon´s diversity index (Welch’s t-test, P<0.05) with higher values observed in samples amplified with HotStarTaq (Fig. 1(b)). Shannon´s diversity index on the samples amplified with HotStarTaq ranged from 7.06 to 9.94 (mean: 8.12 ± 0.97) whereas on the samples amplified with KAPA HiFi the index values ranged from 5.64 to 7.99 (mean: 6.86 ± 0.88).

To assess beta-diversity, we characterised the relative abundance and composition of the *Legionella* phylotypes in the freshwater samples amplified with each DNA polymerase (HotStarTaq and KAPA HiFi) and calculated the Bray-Curtis similarity, using PRIMER (Version 7.0.7) [20], to compare the level of community structure similarity between different samples and between the same sample amplified with the two different DNA polymerases (**Fig. 2**). The term *Legionella* phylotype is used according to what has been previously described [17]. Briefly, NGS sequences were assigned to a species when sequence identity ≥ 97%. The term phylotype englobes the acknowledged *Legionella* species as well as other defined sequence clusters with sequence identity ˂97% to a known species.

**Fig. 2.**
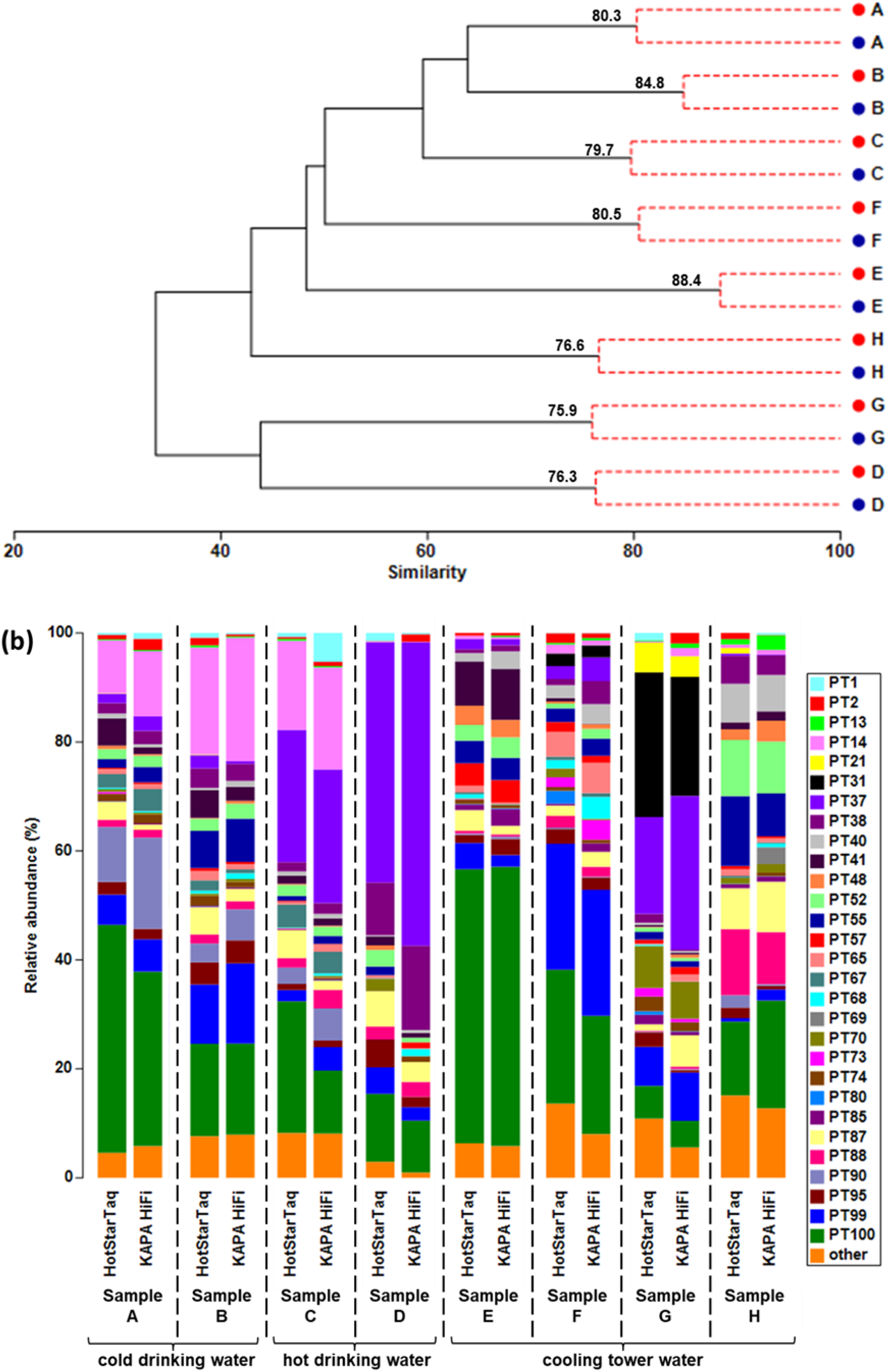
Effect of DNA polymerases on assessment of *Legionella* communities profiling. **(a)** Dendogram showing group-average hierarchical clustering of *Legionella* communities amplified in 8 water samples with both HotStarTaq (blue circle) and KAPA HiFi (red circle) DNA polymerases, using Bray-Curtis similarity. SIMPROF test was performed and red dashed lines in cluster analysis represent groups of samples that do not significantly differ in their *Legionella* community (P>0.05). Values on branches indicate Bray-Curtis similarity index. Sample A, cold drinking water March 2009; sample B, cold drinking water April 2009; sample C, hot water March 2009; sample D, hot water January 2014; sample E, cooling tower water April 2013; sample F, cooling tower water June 2013; sample G, cooling tower water August 2013; sample H, cooling tower water January 2015. **(b)** Bar graph representing relative abundances of *Legionella* phylotypes in 8 water samples amplified with both HotStarTaq and KAPA HiFi DNA polymerase. Phylotypes labelled from 1 to 52 correspond to described *Legionella* species. PT1, L. adelaidensis; PT2, *L. anisa*; PT13, *L. drozanskii*; PT14, *L. dumofii*; PT21, *L. gratiana*; PT31, *L. maceachernii*; PT37, *L. pneumophila*; PT38, *L. quateirensis*; PT40, *L. rowbothamii*; PT41, *L. rubrilucens*; PT48, *L. tucsonensis*; PT52, *L. worsleiensis*. Different freshwater samples are separated by a dashed line.

Based on SIMPROF analysis (Fig. 2(a)) [21], we observed that the difference introduced in the *Legionella* community structure by the use of the two different enzymes was not statistically significant in the sample set used (SIMPROF, P>0.05).

Nonetheless, shifts in the *Legionella* community of every sample studied were observed (Fig. 2(b)), with the similarity levels ranging from 75.9 (Sample G) to 88.4 (Sample E), with an averaged value of 80.3 ± 4.4. Clear clusters for every sample, independently of the type of water, were formed and unambiguously separated the 8 different water samples tested. Furthermore, analysis of the composition of the *Legionella* community in each sample amplified with the two DNA polymerases, revealed a substantial overlap of *Legionella* phylotypes. When the composition of the communities amplified by the two enzymes were compared, 7 out of 8 samples shared more than 85% of the phylotypes (data not shown). Sample D showed the lowest percentage of shared phylotypes (60.1%).

## DISCUSSION

Currently, NGS technologies are widely applied in the characterisation of communities, discovery of novel microorganisms and pathogen detection [22]. For NGS assays to succeed on these fields, a high amplification efficiency during library preparation and target enrichment is of critical importance to downstream sequencing, especially when targeting specific low-abundant taxa in complex environmental samples. The amplification efficiency is influenced by several PCR-related factors, including the choice of DNA polymerase. Our results, despite a better performance of KAPA HiFi, confirmed that the two different polymerases had a very good amplification performance allowing a subsequent appropriate sequencing depth of the target. This was in a certain amount expected due to the hot-start nature of the two enzymes. Plus, our findings highlight that the choice of DNA polymerase, as well as optimisation of the cycling parameters in PCR-based NGS assays, is important to find an appropriate balance between non-specific amplification and yield, in order to maximise detection/sensitivity.

In addition, several studies have reported the impact of different amplification and sequencing-related factors on high-throughput studies of microbial communities and how these might distort diversity estimates and community profiling [23–25]. Yet, only a few studies have evaluated the effects of DNA polymerases on the study of communities [26–29].

Past NGS studies have revealed increasing richness values with increasing sequencing depth [17,30]. Yet, our results showed that the freshwater samples amplified by HotStarTaq DNA polymerase, despite a lower sequencing depth, had significantly higher OTU-based alpha-diversity metrics than KAPA HiFi DNA polymerase, indicating that alpha-diversity measurements are enzyme-dependent, as previously suggested [26–29].

A recent study by Pereira et al. [17], using the same approach, revealed that when comparing the sequence accuracy and quality of the generated NGS *Legionella* libraries, KAPA HiFi had a significantly higher fidelity, with a mean error rate of 0.38% against 0.45% and 65% of error-free sequences against 39%. All data considered, the significant increase in bacterial community richness and diversity, plus the significant higher number of singletons and doubletons observed after amplification of the same 16S rRNA region with HotStarTaq DNA polymerase, is almost certainly linked to a higher generation of erroneous sequences with an enzyme without proofreading activity. This leads to an increased number of OTUs and consequently an artificial overestimation of the richness and diversity estimators [23,31,32]. Similar inflated community richness with polymerases presenting lower fidelity was observed in bacterial libraries by Wu et al. [26] and fungal libraries by Oliver et al. [28]. Yet, conflicting reports by Ahn et al. [27] and Qiu et al. [29] have detected an overestimation of richness with a high-fidelity enzyme instead, attributing it to a higher frequency of chimeric sequences. In our study, as we are targeting a specific taxa, low diversity libraries are generated, not allowing to precisely quantify the potential amount of chimeric sequences in the water samples. However, analysis of *Legionella* mock communities with the same NGS approach has revealed a small representation of these spurious sequences with KAPA HiFi [17].

When comparing beta-diversity metrics after amplification with both DNA polymerases, the impact of the enzymes on the *Legionella* community composition and structure (beta-diversity) was not significant for the samples studied, contrarily to what was indicated by Wu et al. [26], when comparing the DNA polymerases TAKARA ExTaq with PfuUltra II Fusion HS. Though HotStarTaq and KAPAHiFi have different properties, kinetics and slightly different polymerase cocktails were used, the two enzymes seem to have an alike amplification profile of the different *Legionella* phylotypes. However, differences in Bray-Curtis similarity between the samples amplified by two distinct DNA polymerases were observed (mean BC: 80.3 ± 4.4). These discrepancies on *Legionella* community profiling are slightly higher to the ones previously observed between technical replicates (mean BC: 86.8 ± 4.9) [17].

In summary, our data highlights the potential advantageous effects of the use of proof-reading high-fidelity enzymes such as KAPA HiFi on NGS library preparation methodologies. As well, this study emphasises the influence the choice of DNA polymerase has on the characterisation of microbial communities, especially in alpha-diversity metrics, and the critical importance of taking this factor in consideration in comparative studies and in the future use of these high-throughput technologies for pathogen detection and quantification.

## ACKNOWLEDGEMENTS

We gratefully acknowledge Josefin Koch, Marina Pecellin, René Lesnik and Verena Maiberg for their work in environmental sampling and DNA extraction.

## REFERENCES

1. van Dijk EL, Auger H, Jaszczyszyn Y, Thermes C. Ten years of next-generation sequencing technology. Trends Genet. 2014;30(9):418–426.

2. Tan B, Ng C, Nshimyimana JP, Loh LL, Gin KY-H et al. Next-generation sequencing (NGS) for assessment of microbial water quality: current progress, challenges, and future opportunities. Front Microbiol. 2015;6:1027.

3. Wang H, Bédard E, Prévost M, Camper AK, Hill VR et al. Methodological approaches for monitoring opportunistic pathogens in premise plumbing: A review. Water Res. 2017;117:68–86.

4. Forbes JD, Knox NC, Ronholm J, Pagotto F, Reimer A. Metagenomics: The next culture-independent game changer. Front Microbiol. 2017;8:1069.

5. Fields BS, Benson RF, Besser RE. Legionella and Legionnaires’ disease: 25 years of investigation. Clin Microbiol Rev. 2002;15(3):506–526.

6. Pereira RPA, Peplies J, Höfle MG, Brettar I. Bacterial community dynamics in a cooling tower with emphasis on pathogenic bacteria and *Legionella* species using universal and genus-specific deep sequencing. Water Res. 2017;122:363–376.

7. Gargis AS, Kalman L, Berry MW, Bick DP, Dimmock DP et al. Assuring the quality of next-generation sequencing in clinical laboratory practice. Nat Biotechnol. 2012;30(11):1033–1036.

8. Matthijs G, Souche E, Alders M, Corveleyn A, Eck S et al. Guidelines for diagnostic next-generation sequencing. Eur J Hum Genet. 2016;24(1):2–5.

9. Hardwick SA, Deveson IW, Mercer TR. Reference standards for next-generation sequencing. Nat Rev Genet. 2017;18(8):473–484.

10. Eckert KA, Kunkel TA. DNA polymerase fidelity and the polymerase chain reaction. PCR Methods Appl. 1991;1(1):17–24.

11. Chandler DP, Fredrickson JK, Brockman FJ. Effect of PCR template concentration on the composition and distribution of total community 16S rDNA clone libraries. Mol Ecol. 1997;6(5):475–482.

12. Sipos R, Székely AJ, Palatinszky M, Révész S, Márialigeti K et al. Effect of primer mismatch, annealing temperature and PCR cycle number on 16S rRNA gene-targetting bacterial community analysis. FEMS Microbiol Ecol. 2007;60(2):341–350.

13. Brandariz-Fontes C, Camacho-Sanchez M, Vilà C, Vega-Pla JL, Rico C et al. Effect of the enzyme and PCR conditions on the quality of high-throughput DNA sequencing results. Sci Rep. 2015;5:8056.

14. Eichler S, Christen R, Höltje C, Westphal P, Bötel J et al. Composition and dynamics of bacterial communities of a drinking water supply system as assessed by RNA- and DNA-based 16S rRNA gene fingerprinting. Appl Environ Microbiol. 2006;72(3):1858–1872.

15. Henne K, Kahlisch L, Höfle MG, Brettar I. Analysis of structure and composition of bacterial core communities in mature drinking water biofilms and bulk water of a citywide network in Germany. Appl Environ Microbiol. 2012;78(10):3530–3538.

16. Kahlisch L, Henne K, Draheim J, Brettar I, Höfle MG. High-resolution in situ genotyping of Legionella pneumophila populations in drinking water by multiple-locus variable-number tandem-repeat analysis using environmental DNA. Appl Environ Microbiol. 2010;76(18):6186–6195.

17. Pereira RPA, Peplies J, Brettar I, Höfle MG. Development of a genus-specific next generation sequencing approach for sensitive and quantitative determination of the *Legionella* microbiome in freshwater systems. BMC Microbiol. 2017;17:79.

18. Pruesse E, Quast C, Knittel K, Fuchs BM, Ludwig W et al. SILVA: a comprehensive online resource for quality checked and aligned ribosomal RNA sequence data compatible with ARB. Nucleic Acids Res. 2007;35(21):7188–7196.

19. Robertson CE, Harris JK, Wagner BD, Granger D, Browne K et al. Explicet: graphical user interface software for metadata-driven management, analysis and visualization of microbiome data. Bioinformatics. 2013;29(23):3100–3101.

20. Clarke K, Warwick R. Change in marine communities: an approach to statistical analysis and interpretation. 3rd ed. Primer-E Ltd, Plymouth, UK; 2014.

21. Clarke KR. Non-parametric multivariate analyses of changes in community structure. Aust J Ecol. 1993;18(1):117–143.

22. Pallen MJ. Diagnostic metagenomics: potential applications to bacterial, viral and parasitic infections. Parasitology. 2014;141(14):1856–1862.

23. Pinto AJ, Raskin L. PCR biases distort bacterial and archaeal community structure in pyrosequencing datasets. PloS One. 2012;7(8):e43093.

24. Lee CK, Herbold CW, Polson SW, Wommack KE, Williamson SJ et al. Groundtruthing next-gen sequencing for microbial ecology-biases and errors in community structure estimates from PCR amplicon pyrosequencing. PloS One. 2012;7(9):e44224.

25. Kennedy K, Hall MW, Lynch MDJ, Moreno-Hagelsieb G, Neufeld JD. Evaluating bias of Illumina-based bacterial 16S rRNA gene profiles. Appl Environ Microbiol. 2014;80(18):5717–5722.

26. Wu J-Y, Jiang X-T, Jiang Y-X, Lu S-Y, Zou F et al. Effects of polymerase, template dilution and cycle number on PCR based 16S rRNA diversity analysis using the deep sequencing method. BMC Microbiol. 2010;10:255.

27. Ahn J-H, Kim B-Y, Song J, Weon H-Y. Effects of PCR cycle number and DNA polymerase type on the 16S rRNA gene pyrosequencing analysis of bacterial communities. J Microbiol. 2012;50(6):1071–1074.

28. Oliver AK, Brown SP, Callaham Jr. MA, Jumpponen A. Polymerase matters: non-proofreading enzymes inflate fungal community richness estimates by up to 15%. Fungal Ecol. 2015;15:86–9.

29. Qiu X, Wu L, Huang H, McDonel PE, Palumbo AV et al. Evaluation of PCR-generated chimeras, mutations, and heteroduplexes with 16S rRNA Gene-Based Cloning. Appl Environ Microbiol. 2001;67(2):880–7.

30. Claesson MJ, Wang Q, O’Sullivan O, Greene-Diniz R, Cole JR et al. Comparison of two next-generation sequencing technologies for resolving highly complex microbiota composition using tandem variable 16S rRNA gene regions. Nucleic Acids Res. 2010;38(22):e200.

31. Kunin V, Engelbrektson A, Ochman H, Hugenholtz P. Wrinkles in the rare biosphere: pyrosequencing errors can lead to artificial inflation of diversity estimates. Environ Microbiol. 2010;12(1):118–23.

32. Reeder J, Knight R. The ‘rare biosphere’: a reality check. Nat Methods. 2009;6(9):636–7.

